# How antiporters exchange substrates across the cell membrane? An atomic-level description of the complete exchange cycle in NarK

**DOI:** 10.1101/2020.06.02.130773

**Authors:** Jiangyan Feng, Balaji Selvam, Diwakar Shukla

## Abstract

Major facilitator superfamily (MFS) proteins operate via three different mechanisms: uniport, symport, and antiport. Despite extensive investigations, molecular understanding of antiporters is less advanced than other transporters due to the complex coupling between two substrates and the lack of distinct structures. We employ extensive (~300 *μ*s) all-atom molecular dynamics simulations to dissect the complete substrate exchange cycle of the bacterial 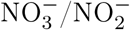 antiporter, NarK. We show that paired basic residues in the binding site prevent the closure of unbound protein and ensure the exchange of two substrates. Conformational transition only occurs in the presence of substrate, which weakens the electrostatic repulsion and stabilizes the transporter by ~1.5 ± 0.1 kcal/mol. Furthermore, we propose a state-dependent substrate exchange model, in which the relative spacing between the paired basic residues determines whether 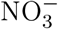 and 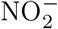 bind simultaneously or sequentially. Overall, this work presents a general working model for the antiport mechanism within MFS family.

## Introduction

Major facilitator superfamily (MFS) transporters move a wide spectrum of biologically relevant substrates (nutrients, drugs, waste, etc.) across cell membranes.^1–4^ This superfamily is ubiquitous in all kingdoms of life and represents the largest and most diverse group of secondary active transporters.^5^ Malfunction of MFS proteins has been associated with a multitude of diseases, such as cancer, type 2 diabetes, and Alzheimer’s disease.^6–8^ In plants, MFS transporters mediate macronutrients (C, N, and P) uptake and extrusion of deleterious compounds.^9–11^ Owing to their physiological and pathophysiological significance, MFS proteins have been popular targets for structural and mechanistic investigations. MFS transporters can be classified into three types depending on their substrate-transport mechanism: uniporters, symporters, and anitporters.^12^ Uniporters transport a single species of substrate across the membrane. Symporters translocate two or more substrates in the same direction. Antiporters transport a substrate and a co-substrate in opposite directions. Our understanding of transport mechanism is well advanced for symporters and uniporters, exemplified by the *Escherichia coli* lactose/H^+^ symporter (LacY).^13–15^ By contrast, molecular insights are less advanced for antiporters due to the lack of crystal structures representing different stages of the transport cycle. ^5^ Among all the MFS proteins with known structures, most of them are symporters and the outward-facing conformation of MFS antiporters has yet to be captured.^16^ Recently, computational studies of a few antiporters have advanced our understanding of antiport mechanism.^5,17–20^ However, these studies were only able to simulate the transport of one substrate in the transport cycle, even with enhanced sampling techniques and state-of-the-art supercomputing.

Many important structural and mechanistic aspects of antiport mechanism remain elusive: (1) Can antiporters be distinguished from other transporters (uniporters and symporters) based on their protein architecture and conformational transition mechanism? (2) How is the empty transporter prevented from changing conformations in antiporters? (3) How does an antiporter distinguish and switch two cargos? (4) How is substrate binding coupled with global conformational changes? (5) Which residues regulate substrate recognition, binding, and release?

NarK represents a convenient model system for studying the functions of MFS antiporters because it has been characterized in a few conformations.^21,22^ NarK is a 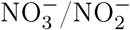 antiporter from *Escherichia coli*, coupling an outward flow of internal 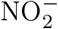 to the uptake of 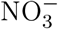 into cell.^22^ It belongs to the 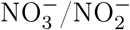 porter (NNP) family of the MFS. NNP proteins mediate the high-affinity translocation of 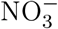 and 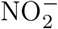 in both prokarytoes and eukaryotes (*i.e.* archaea, bacteria, fungi, yeast, algae, and plants).^23^ 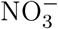 is the most important source of mineral nitrogen for plant and 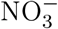 availability greatly limits crop yields.^24^ Plant NNP members (e.g. *Arabidopsis thaliana* NRT2.1 and NRT2.2) account for 80% of high-affinity 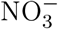 uptake, rendering them essential under low nitrogen conditions. ^9^ A detailed understanding of the structure, transport mechanism, and conformational changes of NNP transporters would therefore substantially guide the engineering of these transporters to enhance agricultural productivity. Recent crystal structures revealed that NarK adopts a canonical MFS fold consisting of 12 transmembrane (TM) helices. These helices are organized into two structurally similar domains, the N-domain (TM1-TM6) and the C-domain (TM7-TM12). Membrane transporters generally work by an alternating-access mechanism.^25–27^ Switching among outward-facing (OF), occluded (OC), and inward-facing (IF) states alternatively expose the substrate binding site to either side of the membrane.

Although static snapshots of X-ray crystallography are critical, they are insufficient to explain the mechanistic details of such dynamic transitions and their coupling to chemical events supplying the energy. Extensive sets of molecular dynamics (MD) simulations^28–30^ have been successfully combined with Markov state models (MSMs)^31,32^ to reveal the dynamics and conformational transitions of various membrane proteins including transporters.^10,11,33–37^ MSMs stitch massive parallel short MD trajectories together to build a kinetic network model that describes long timescale protein dynamics. ^31,32^

This work explores the structural and mechanistic principles that characterize MFS antiporters through the study of a bacterial 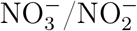 antiporter, NarK. By combining extensive (~300 *μ*s) unbiased all-atom MD simulations and MSMs, we explored the complete substrate exchange cycle for NarK and characterized the underlying free energy landscapes. Simulation results suggest that NarK adopts the conserved MFS fold and follows the common rocker-switch mechanism, governed by helix bending around highly conserved glycine residues. What distinguishes NarK antiporter from symporters is the inaccessible energy barrier between IF and OF states without a bound substrate. Two highly conserved and positively charged arginine residues (R89 and R305) in the central binding site restrict the closure of unbound protein, thereby ensuring the exchange of 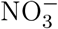 and 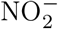. Substrate binding (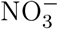 or 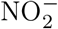) is required to weaken the electrostatic repulsion and drive the conformational switch. We further identified key residues involved in substrate recognition, binding, exchange, and translocation, and provided a detailed model of the complete exchange process. This work provides important information both for understanding 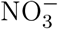 uptake by NNP transporters and for elucidating the antiport mechanism within MFS family.

## Results and Discussion

### NarK follows the canonical rocker-switch mechanism

All-atom MD simulations of NarK were initiated from the 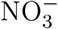-bound occluded crystal structure (PDB ID: 4U4W^22^). In all these simulations, the transporter was inserted into an explicit lipid bilayer and solvated in a box with dimension of 76Å*76Å*102Å containing TIP3P water molecules. The simulations were carried out under isothermal-isobaric (NPT) condition (300K and 1 atm). With a total of ~300 *μ*s unbiased MD simulations, the entire transport cycle of NarK and all the functionally relevant states were characterized. Markov state model (MSM) is a popular technique for extracting kinetic information of protein dynamics from MD simulation data.^31,32^ All the simulation data were used to construct an MSM and the MSM hyper-parameters were selected systematically using a genetic algorithm technique.^38^ The MSM estimation reweighs the MD trajectories such that the equilibrium kinetics and distribution among sampled configurations can be recovered. Simulation and MSM construction details are summarized in Method Details. Because only a finite quantity of simulation data can be obtained to construct the MSM model, the properties computed from the MSM will be statistically uncertain. Using a Gibbs sampling procedure, Bayesian MSMs construct a sample of reversible transition matrices and quantify the statistical uncertainties for all observables derived from MSM.^39,40^ To estimate the uncertainties, a Bayesian MSM^39,40^ was estimated with 100 samples and 95% confidence interval using PyEMMA2.5.7^41^ (see Quantification and Statistical Analysis for details). In the crystal structure (PDB ID: 4U4W^22^), Ser56 (TM1) hydrogen bonds with Ala275 (TM7) at the extracellular side and closes the pore tunnel. The hydrophobic interactions between Met151 (TM4) and Phe370 (TM10) act as an intracellular gate. We projected simulation data onto these two metrics as the opening and closure of the pore channel strictly determines the specific state such as IF, OC, and OF.^33^

Free energy landscape reveals the thermodynamic and kinetic information about the antiporter transport mechanism (Fig.1A). The canonical L-shaped free energy landscape displays three distinct free energy basins corresponding to IF, OC, and OF states, in line with the rocker-switch mechanism. The three crystal structures of NarK were labeled in the free energy landscape. Simulations predict that these crystal structures^22^ are all from the most stable basin of the transporter, the IF state. The measurement of extracellular and intracellular distances shows that the helices of the N and C domains are ~4.7 and ~13.2 Å apart in the crystallized IF state (PDB ID: 4U4T, 4U4V^22^). However, the free energy landscape suggests that the intracellular distance between TM4 and TM10 can increase up to ~15.0 Å for the energetically accessible IF state (Fig.1A and Supplementary Fig.1A, B). As TM4 and TM10 move towards each other, NarK adopts an OC state with intracellular distance reduced to ~9.0 Å while extracellular distance increased to ~7.5 Å. According to these metrics at least, the previously reported OC state (PDB ID: 4U4W^22^) may actually represent a partially occluded IF state. The intracellular side can be further closed as the intracellular distance decreasing from ~11.5 to ~9.0 Å (Fig.1A and Supplementary Fig.1C, D). Finally, the extracellular distance increases to ~12.0 Å and the antiporter adopts the OF state. The free energy barrier for the transition from IF to OC state is ~1.5 ± 0.1 kcal/mol and for the subsequent transition to OF state is ~1.0 ± 0.1 kcal/mol. The total free energy barrier for one complete cycle of NarK from IF to OF state is ~2.5 ± 0.1 kcal/mol. The relatively low energy barrier suggests that NarK can easily interconvert between IF, OC, and OF states during the translocation process. This observation is consistent with the high-affinity uptake of 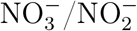 in NarK.^23,42^

**Figure 1:**
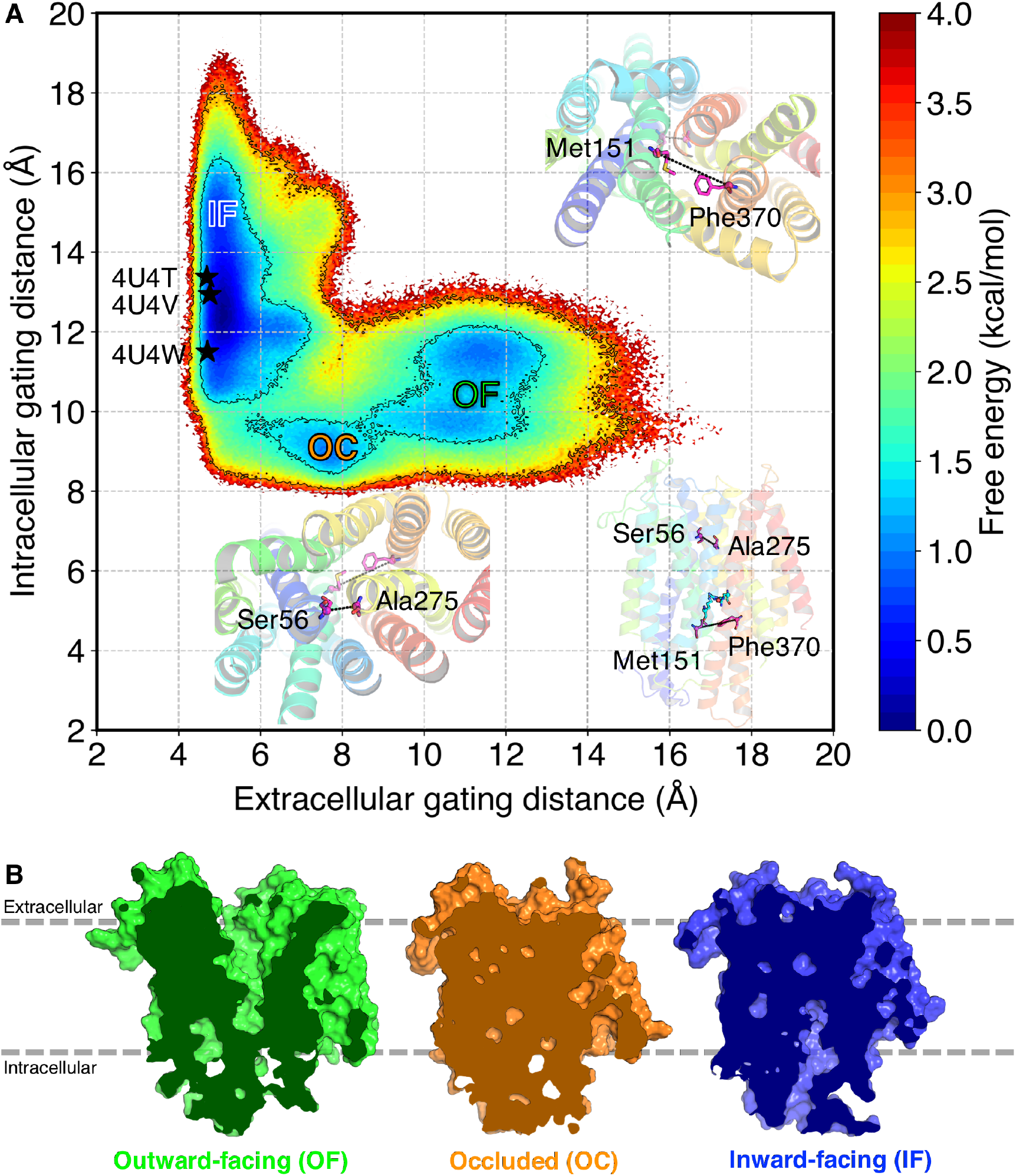
Alternating access cycle of NarK. A. MSM-weighted free energy landscape plot of NarK. The ~300 *μ*s MD simulation data were projected onto two-dimensional space of Ser56C*α* (TM1) - Ala275C*α* (TM7) and Met151C*α* (TM4) - Phe370C*α* (TM10) distances. Three crystal structures were labelled as black stars. B. Cross-sections of surface representations for the representative structures, sampled randomly from the corresponding OF, OC, and IF states. The difference between the intracellular and extracellular gating distance (Δ *d*) was used as the selection criteria for the OF (Δ *d* ~ − 1.3 ± 0.01 Å), OC (Δ *d* ~ 3.3 ± 0.01) Å, and IF (Δ *d* ~ 10.8 ± 0.01 Å) states. The extracellular gating distance is 12.3 Å, 5.1 Å, and 4.4 Å for the shown OF, OC, and IF structures. The intracellular distance is 11.0 Å, 8.6 Å, and 15.2 Å for the shown OF, OC, and IF structures. The structures were viewed from the membrane plane.

To gain structural insights into the free energy landscape, three representative structures were randomly sampled from the corresponding OF, OC, and IF states (Fig.1B, coordinate files have been deposited to Github: https://github.com/ShuklaGroup/NarK_Structure_2021_Files). The difference between the intracellular and extracellular gating distance (Δ*d*) was used for selecting OF (Δ*d* ~ −1.3 ± 0.01 Å), OC (Δ*d* ~ 3.3 ± 0.01) Å, and IF (Δ*d* ~ 10.8 ± 0.01 Å) conformations. The molecular surfaces for these structures suggest that the substrate translocation pathway is the pore saddled by the N- and C-terminal domains. N- and C-domains change their relative positions to alternatively expose the substrate binding site to opposite sides of the membrane. These structural features reinforce the notion that NarK follows the rocker-switch mechanism. ^27^ The simulated IF and OC structures show good agreement with experimental NarK structures (PDB ID: 4U4V, 4U4T, 4U4W^22^) (Supplementary Fig.1). The most pronounced differences are in the opening of intracellular gating for both IF and OC states. The opening at the intracellular side in simulated structures is wider for IF state (~2.0 Å) and narrower in OC state (~2.5 Å). The predicted OF structure was compared to a fucose/H^+^ symporter (FucP, PDB ID: 3O7Q^43^) due to the absence of OF structures in MFS antiporters (Supplementary Fig.1E, F). The OF conformation superimposes well on the OF structure of FucP and exhibits similar opening at the extracellular side (Supplementary Fig.1E, F). These findings suggest that the rocker-switch mechanism is shared among all MFS proteins, irrespective of their particular functions as a uniporter, symporter, or antiporter. Therefore, the diversity of MFS transporter functions is a result of changes in a few residues in the binding pocket and translocation pathway.

### Helix bending drives opening and closing of intracellular and extracellular gates

To investigate the conformational changes underlying the rocker-switch type movement, we computed root-mean-square fluctuation (RMSF) of each C*α* atom in NarK during IF to OC and OC to OF transition (Fig.2A-F, Supplementary Fig.2). The intracellular tips of TM5, TM10, and TM11 show higher flexibility (~2.5 Å) during the IF to OC transition (Fig.2E-F). The extracellular tips of TM1, TM2, and TM7 undergo significant rearrangements (~2.5 Å) during the transition from OC to OF state (Fig.2C-D). The other helices (TM3, TM4, TM6, TM8, TM9, and TM12) represent a stable structural element that displays limited conformational changes (~1.0 Å) during the transport cycle. Previous structural analysis hypothesized the N-bundle remains rigid while TM7, TM10, and TM11 helices of the C-bundle move towards the N-bundle to close the intracellular vestibule. ^22^ In simulations, both the N-(TM5) and C-domains (TM10 and TM11) move towards each other with a higher magnitude (~1.0 Å higher) for the C-domain to close the intracellular side. It was also suggested that TM7 is involved in the closure of intracellular gate.^22^ By contrast, the extracellular tip of TM7 exhibits large fluctuations (~2.5 Å) during OC to OF transition whereas the motions in the intracellular tip are negligible (~1.0 Å) during the entire transport cycle (Fig.2 C, D, G). This strongly indicates that TM7 is critical for the opening of the extracellular instead of intracellular gate. The simulations also shed light on the conformational transitions from OC to OF states, which were previously unknown due to the lack of an OF structure. The extracellular half of TM1 and TM2 from the N-domain, and TM7 of the C-domain move apart to open the extracellular vestibule.

**Figure 2:**
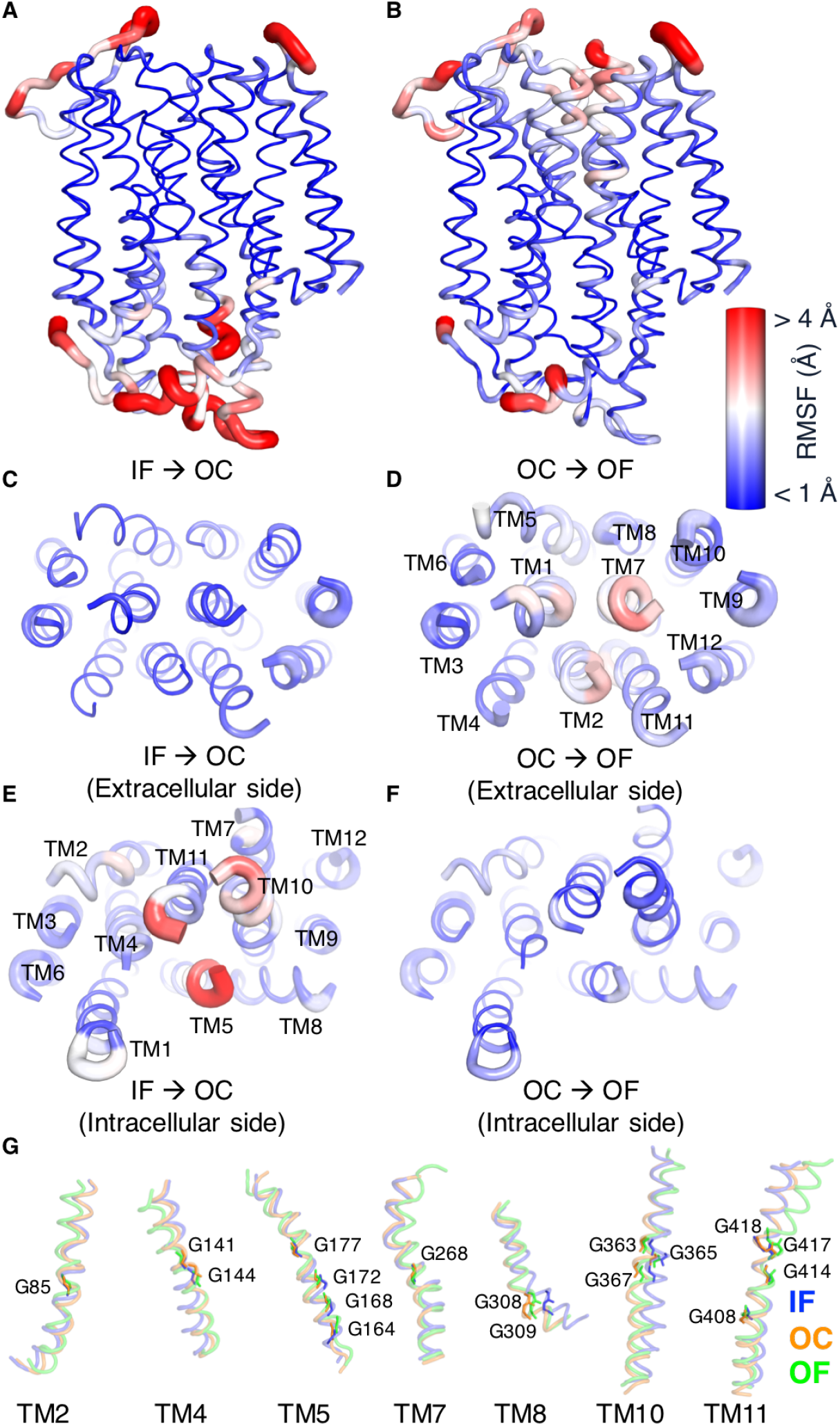
Global conformational changes. Root-mean-square fluctuation (RMSF) data were mapped onto the predicted IF structure of NarK for the IF to OC (A, C, E), and OC to OF transitions (B, D, F). Blue and red correspond to low (< 1 Å) to high (> 4 Å) RMSF values. Tube thickness corresponds to the RMSF value of each residue. Close-up views from the extracellular (C, D) and intracellular (E, F) side with transmembrane helices (TMs) labelled. G. Structural comparisons among IF, OC, and OF states. The superimposition was performed with Pymol^49^ and for the helices containing conserved glycine residues: TM2, TM4, TM5, TM7, TM8, TM10, and TM11. The helices were shown in tube representations and colored based on the conformational states. Relevant glycine residues were labelled and shown as colored sticks.

The RMSF data illustrate that only helix tips show significantly higher flexibility over the entire transport cycle from IF to OC and OC to OF (Fig.2A-F, Supplementary Fig.2). These results suggest that the conformational transition between IF and OF is governed by the internal bending and straightening motion of the helices (TM1, TM2, TM5, TM7, TM10, and TM11), rather than the rigid body tilting movement of two bundles. The bending of TM10 and TM11 was also observed in the previous structural studies of NarK,^21,22^ its closest homolog NarU, ^42^ and many other MFS family proteins,^43–47^ implying a significance of flexibility of TM10 and TM11 within MFS family. An important feature of NNP family transporters is the presence of highly conserved glycine residues, which constitute the inner core of many TMs (TM2, TM4, TM5, TM7, TM8, TM10, and TM11).^21,22,42,48^ It is interesting to note that large structural fluctuations in helices are located at or close to the conserved glycine residues (Fig.2G). These highly conserved glycine residues serve as pivot points for helix bending. The significance of glycine is underscored by previous mutations of glycine to alanine that abolish the 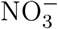 uptake activity.^22^ Together, our simulations revealed that the helix bending around highly conserved glycine residues governs the overall conformational change in NarK, and that these highly conserved glycine residues may support the structural flexibility in all members of the NNP family.

### Hydrophobic and polar interactions lock the transport pathway

To identify the structural features that support the helix bending motions, we computed the key interactions within N- and C-terminal domains in three different conformational states (IF, OC, and OF). 10 sets of 1,000 MD configurations were randomly selected for each state according to the difference between the intracellular and extracellular gating distance (Δ*d*): OF (Δ*d* ~−1.3 ± 0.01 Å), OC (Δ*d* ~ 3.3 ± 0.01) Å, and IF (Δ*d* ~ 10.8 ± 0.01 Å). By comparing the average contact frequency in different states, we identified six layers of hydrophobic and polar interactions along the transport pathway (Fig.3). Many of these interactions were not reported in the previous studies which rely on the structural overlays of IF and partially occluded IF state.^21,22^ Simulation data suggest that helix bending regulates the disruption and formation of these interactions and thus controls the opening and closing of the gates.

**Figure 3:**
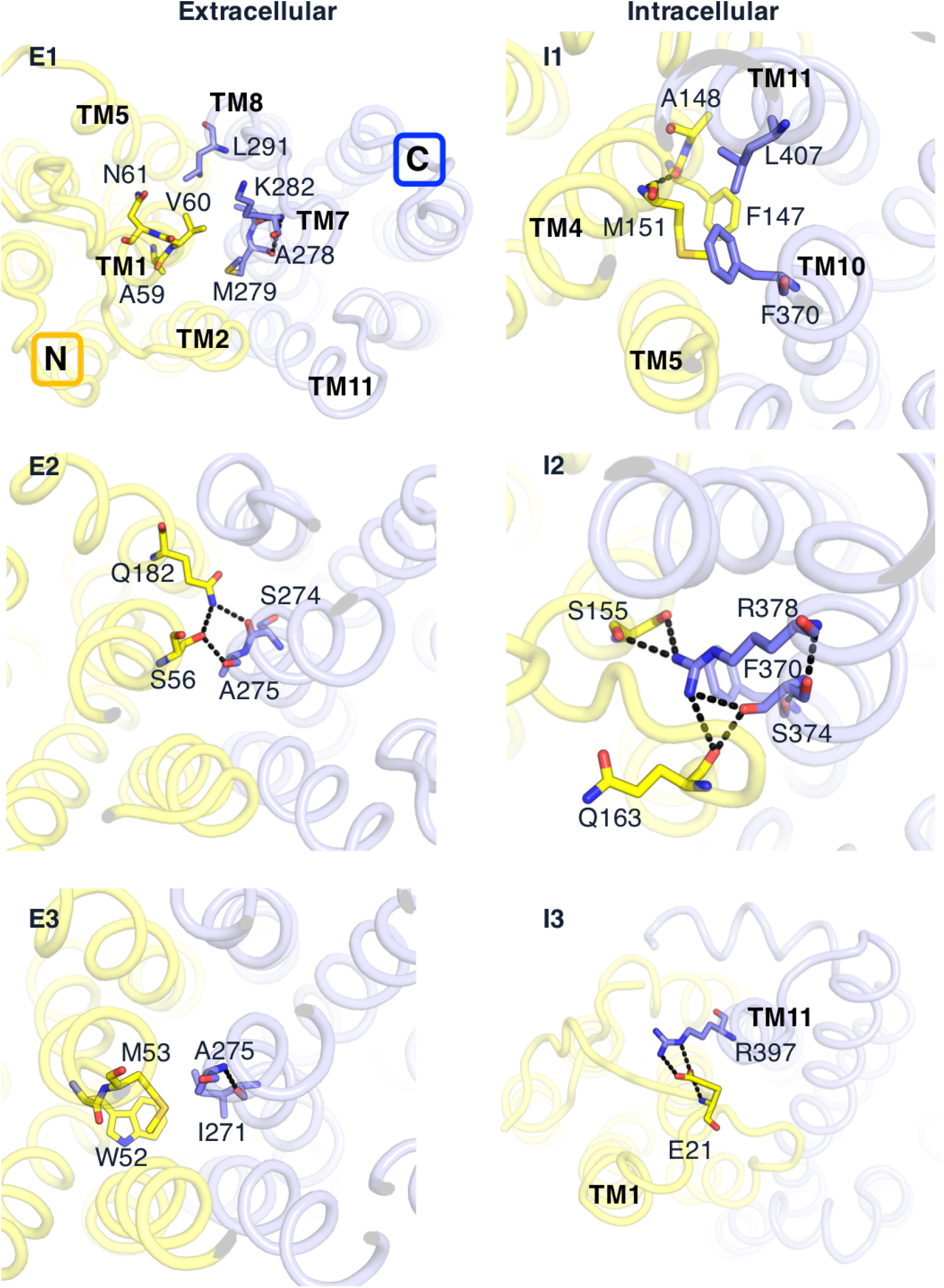
Extensive hydrophobic and polar interactions along the transport pathway. The three left panels represent the close-up views of the extracellular gating interactions seen from the extracellular side. The three right panels show the close-up views of the intracellular gating interactions seen from the intracellular side. N- and C-domain helices were colored yellow and light blue, respectively. The residues involved in each interaction layer were depicted by stick models and colored following the same scheme.

The exit of substrates to the extracellular side is blocked by a number of gating residues mainly in TM1 and TM7 (Fig.3 E1-3). At the extracellular side, residues A59 (TM1), V60 (TM1), N61 (TM1), A278 (TM7), M279 (TM7), K282 (TM7), and L291 (TM8) constitute the hydrophobic layer E1 (Fig.3 E1). All of these residues are conserved in NarK’s closest homolog (NarU), except A59, suggesting a similar hydrophobic layer in NarU. ^42^ In layer E2, S56 (TM1), Q182 (TM5), S274 (TM7), and A275 (TM7) form an extensive network of hydrogen bonds, facilitating the close packing of gating helices (Fig.3 E2). This polar interaction layer was also reported in NarU and alanine substitutions of S54 and Q180 (equivalent to S56 and Q182 in NarK) disrupted the binding reactions. ^42^ The hydrophobic residues in layer E3 (W52 (TM1), M53 (TM1), I271 (TM7), and A275 (TM7)) mediate extensive van der Waals contacts and stabilize the extracellular gate (Fig.3 E3). W52 is invariant among all the prokaryotic NNP family transporters. W50A mutation in NarU (corresponding to W52 in NarK) completely abolished binding to both 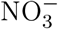 and 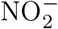, implying a critical gating role in all prokaryotic NNP transporters. The extracellular gating residues are clustered in TM1 and TM7. Therefore, the bending of TM1, TM2, and TM7 can effectively control whether the extracellular gate is open or closed during the transition between the OF and OC states.

Consistent with experimental structure (PDB ID: 4U4W^22^), the intracellular gate consists of three layers of interactions (Fig.3 I1-3). In layer I1 (immediately beneath the substrate-binding pocket), residues F147, A148, and M151 of TM5 hydrophobically interact with F370 of TM10, and L407 of TM11 (Fig.3 I1). F147 and F370 are invariant among all NarK homologs in bacteria and eukaryotes. Previous functional analysis of NarK and its closest homologue (NarU) further supports the functional relevance.^22,42^ The F147A mutant shows decreased 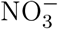-transport activity in NarK^22^ and F367A mutation in NarU^42^ (corresponding to F370 in NarK) abrogated binding to both 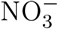 and 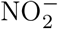. In the previously reported layer I2,^22^ S155 in TM4 hydrogen bonds with A400 in TM11, and F158 and K160 in TM5 hydrogen bond with R378 in TM10. In simulations, however, the S155-A400 and R378-F158-K160 interactions are formed in both IF and OF states (Fig.3 I2). The S155-A400 interaction is observed in around 88% and 93% of IF and OF states. The R378-F158-K160 interaction occurs approximately in 75% and 100% of structures adopting the IF and OF conformations. These interactions are therefore not crucial for the conformational switch from IF to OF states. Simulations suggest that S155(TM4) and Q163 (TM5) form an extensive hydrogen bond network with S374 and R378 from TM10 (Fig.3 I2). In the partially occluded IF state of NarK (PDB ID: 4U4W^22^), Q163 on unbent TM5 points away from the central pathway and thus the extensive polar interactions of layer I2 are missing (Supplementary Fig.3). The bending of TM5 and TM10 is crucial to form the polar interaction layer I2, thereby facilitating a much tighter packing between N and C bundles (Fig.3 I2). Near the intracellular surface, E21 of the N-terminal loop forms salt bridge with R397 on TM11 and blocks the access to the substrate binding site from the cytoplasm (Fig.3 I3). Upon the outward bending of TM11, E21 changes its salt bridge partner from R397 to K160 and allows the opening of the inside gate.

### Substrate unoccupied transporter restricts conformational changes in an antiporter

Given the common conformational transition mechanism, how can uniporters, symporters, and antiporters be distinguished from each other? A major difference between a symporter and an anitporter is whether an empty transporter changes conformation from IF to OF: symporters do, but antiporters do not.^50^ How is such a mechanistic scheme implemented in an antiporter? We projected all the simulation data onto a two-dimensional free energy landscape to understand the whole 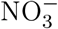 and 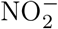 exchange cycle (Fig.4, Supplementary Fig.4). To monitor the conformational transitions from IF to OF state, the difference between the intracellular and extracellular gating distances was selected as x coordinate. Previous mutational analysis demonstrates that R89 and R305 directly bind substrates.^22^ The difference between the closest distances from 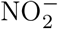 and 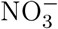 to the center of R89 and R305 was selected as y coordinate to track the substate binding mode.

**Figure 4:**
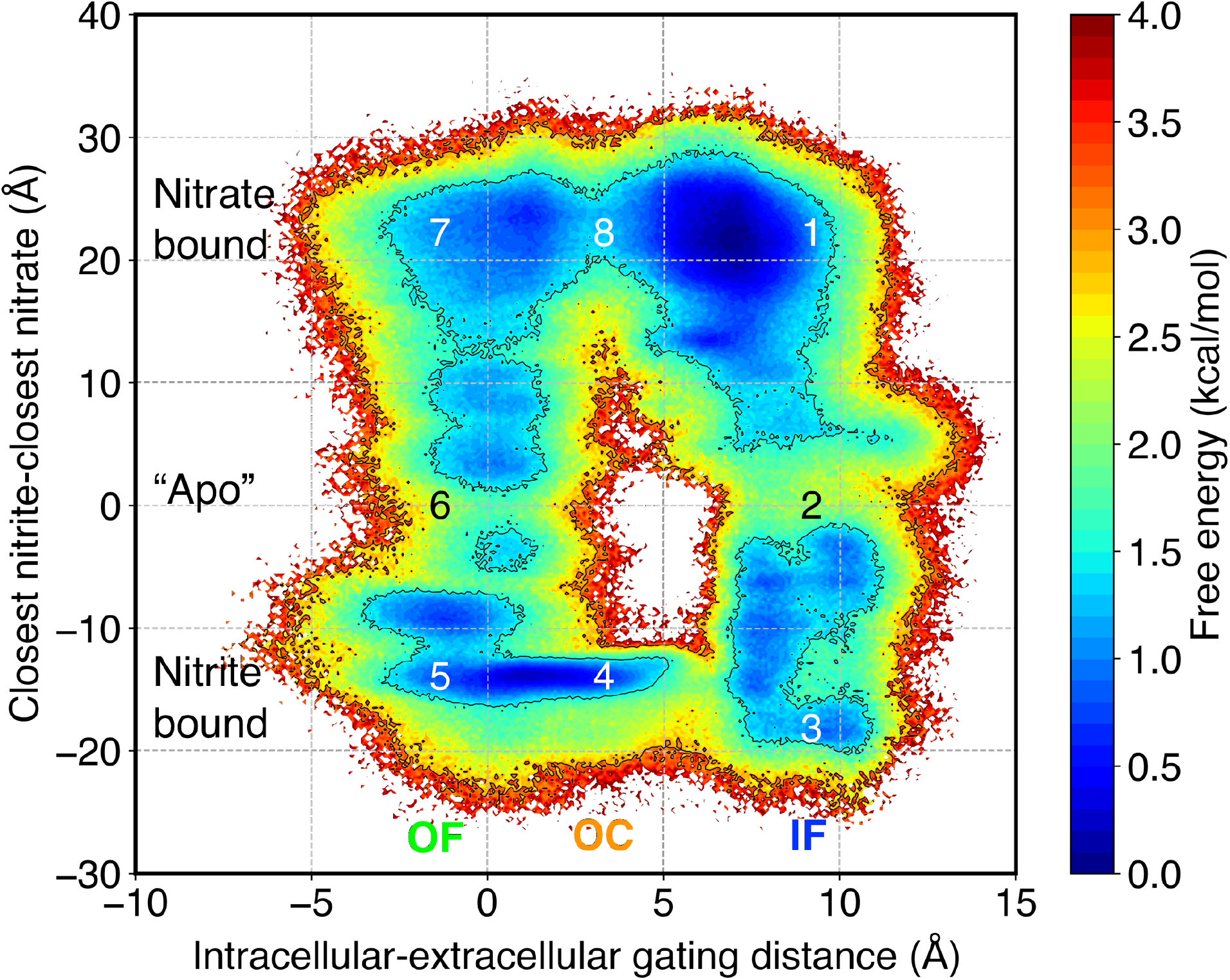
MSM-weighted free energy landscape of the entire 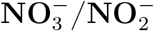 exchange cycle. The difference between the intracellular and extracellular gating distances (x) differentiates IF, OC, and OF states. The difference between the closest distances from 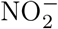 and 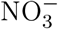 to the center of R89 and R305 (y) shows different substrate binding conditions. The conformational states were depicted as (1) 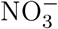-bound IF (x = 9.2 Å, y = 22.0 Å), (2) *“apo”* IF (x = 9.2 Å, y = −0.2 Å), (3) 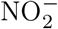-bound IF (x = 9.2 Å, y = −18.4 Å), (4) 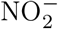-bound OC (x = 3.3 Å, y = −14.2 Å), (5) 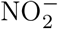-bound OF (x = −1.3 Å, y = −14.2 Å), (6) *“apo"* OF (x = −1.3 Å, y = −0.2 Å), (7) 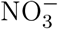-bound OF (x = −1.3 Å, y = 22.0 Å), and (8) 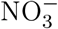-bound OC (x = 3.3 Å, y = 22.0 Å).

The free energy landscape exhibits eight distinct conformational states constituting the whole 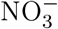 and 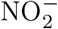 exchange cycle (Fig.4). The white region in the center implies an inaccessible energy barrier between IF and OF states without a bound substate. The high energy barrier excludes the alternation between the OF and IF states without the aid of a substrate, thereby implementing the strict 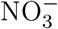 and 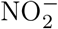 exchanger function. Substrate binding is required in an antiporter to promote the conformational switch. While the substrate bound states correspond to the energy minima, substrate unbound states are located in the relatively higher energy regions (~2.0 ± 0.1 kcal/mol). Substate binding lowers the energy by ~1.5 ± 0.1 kcal/mol in both IF and OF states and enhances probability for the conformational switching between them.

Moreover, the free energy landscape enables the quantitative comparison between the 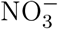 and 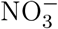 transport dynamics. Although 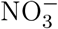 and 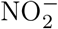 are equal in charge and differ merely by one oxygen atom, pronounced differences between them were found in the energy barrier between IF and OC states. The free energy barrier for the transition between OC and IF states is ~1.0 ± 0.1 kcal/mol with bound 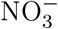. When 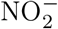 is bound, OC and IF states are separated by a barrier of ~2.5 ± 0.1 kcal/mol. According to the Arrhenius equation,^51^ the ~1.5 ± 0.1 kcal/mol energy difference corresponds to ~12-fold binding affinity difference between 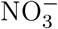 and 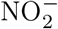. This is consistent with the 10-fold higher binding affinity for 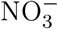 (~33 *μ*M) over 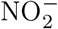 (~373 *μ*M) in NarK’s closest homolog, NarU. ^42^ The computational analyses suggest that 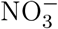 favors the transition from OC to IF states and thus enhances the transport activity.

### 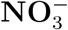 and 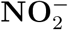 switch differently in IF and OF states

One intriguing aspect of MFS antiporters as a whole is their capability to differentiate and exchange two similar substrates.^5^ To gain structural insights into this capability, we compared eight representative structures randomly sampled from the center of corresponding metastable states in Fig.4 (coordinate files have been deposited to Github: https://github.com/ShuklaGroup/NarK_Structure_2021_Files). Each free energy basin corresponds to a distinct substrate binding mode. In agreement with previous crystal structures,^22^ 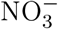 and 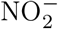 bind to NarK at exactly the same position (Fig.5). This binding pocket is formed by the side chains of highly conserved R89, G144, F147, F49, G171, N175, R305, S366, F267, Y263, and S411 (Fig.5B). The functional significance of these residues has also been confirmed by mutational analysis directly performed on NarK.^22^ Two conserved and positively charged residues R89 from TM2 and R305 from TM8 directly bind to substrates.^22,52^ Mutation of either arginine into lysine decreases transport activity of NarK. Without bound substrate, the electrostatic repulsion between R89 and R305 prevents the closure of the pore channel and thus implements the strict substrate exchange function in NarK. Paired basic residues are also essential for substrate binding in several other MFS antiporters such as *Escherichia coli* sn-glycerol-3-phosphate transporter (GlpT),^53^ *Escherichia coli* Hexose-6-phosphate:phosphate antiporter (UhpT),^54,55^ and the oxalate:formate transporter from *Oxalobacter formigenes* (OxlT).^56,57^ This suggests that paired basic residues may be the structural determinant for the exchange function in antiporters transporting anionic substrates. There is a single binding pocket in NarK for two different substrates. The question is how NarK distinguishes between two similar cargo molecules such as 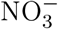 and 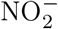. The differences between 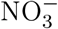 and 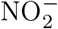 binding are strikingly clear in the OC state (state 4 and 8 in Fig.5B). The additional oxygen atom of 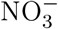 allows the formation of additional hydrogen bonds with R89 and R305 and connects the two halves of NarK tightly. This may accelerate the OC to IF transition and lead to ~1.5 ± 0.1 kcal/mol difference in Fig.4.

**Figure 5:**
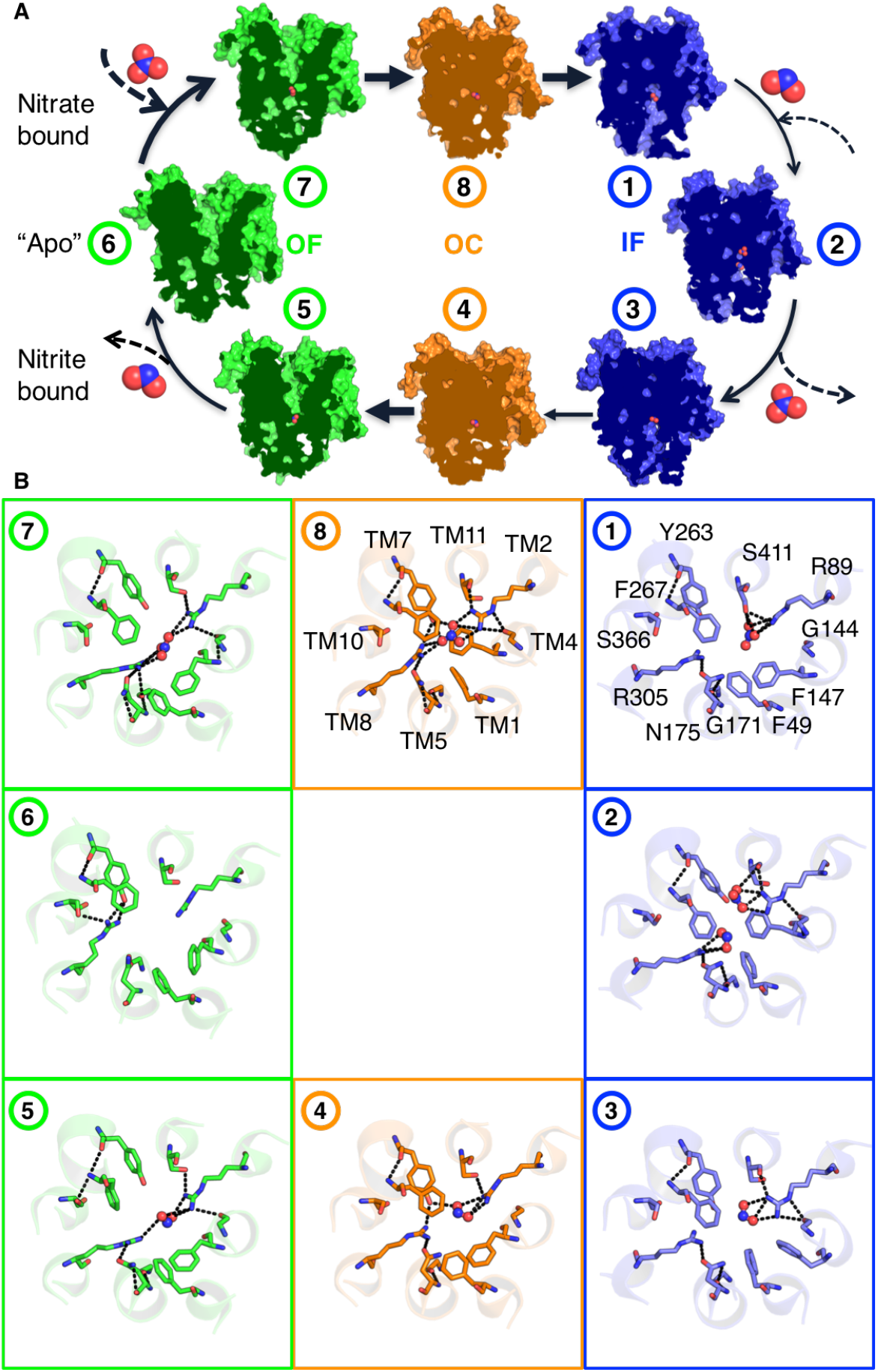
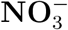 and 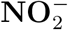 switching mechanism. A. Cross-section of surface representations for the representative structures from the intermediate states along the exchange cycle. The shown conformations were randomly sampled from the center of the eight metastable states in Fig.4: (1) 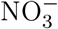-bound IF (x = 9.2 Å, y = 22.0 Å), (2) *“apo"* IF (x = 9.2 Å, y = −0.2 Å), (3) 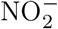-bound IF (x = 9.2 Å, y = −18.4 Å), (4) 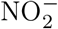-bound OC (x = 3.3 Å, y = −14.2 Å), (5) 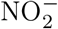-bound OF (x = −1.3 Å, y = −14.2 Å), (6) *“apo"* OF (x = −1.3 Å, y = −0.2 Å), (7) 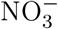-bound OF (x = −1.3 Å, y = 22.0 Å), and (8) 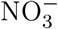-bound OC (x = 3.3 Å, y = 22.0 Å), where x represents the difference between the intracellular and extracellular gating distances and y represents the difference between the closest distances from 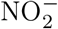 and 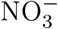 to the center of R89 and R305. The bound substrates (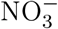 and 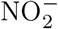) were shown in ball and sticks. B. Close-up view of the binding sites visualizing the residues coordinating binding of 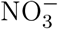 and 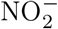. The dotted black lines represent hydrogen bonds among residues. Substrates and binding pocket residues were represented in ball-and-stick and sticks, respectively.

The next question is how antiporter switches the substrates. A common hypothesis for exchangers is the “ping-pong" mechanism.^58^ It assumes that there is only one binding site and that the protein never binds both substrates simultaneously. The antiporter must export a substrate first to import a second substrate. But NarK exhibits state-dependent substrate exchange behavior. When NarK adopts IF state (state 2 in Fig.5B), the binding of both substates is preferred (~65 %) over the unbound conformation (~35 %) (Supplementary Fig.5A). In comparison, ~90 % of structures adopt an unbound conformation in the OF state (state 6 in Fig.5B) (Supplementary Fig.5B). This is likely due to the relative spacing between R89 and R305. In the IF state, the shortest distance between R89 and R305 side chains is ~9.0 Å. R89 and R305 could not simultaneously coordinate the small anion (~1.96 Å for 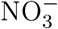) (state 1, 3 in Fig.5B). During the IF to OF transition, helix bending moves R305 and R89 closer to each other by ~2.0 Å, which allows both R89 and R305 to form hydrogen bonds with the substrate (state 5, 7 in Fig.5B). From these data, we propose a state-dependent exchange model. In the IF state, 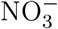 and 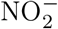 simultaneously bind to NarK. The electrostatic repulsion between the two anions facilitates the release of one substrate and exchanges the substrates. In OF state, the first substrate leaves the binding site and then the second substrate comes in and binds the protein.

### Y263 and R305 couple substrate binding and conformational changes

A fundamental question in the study of MFS proteins is how can local substrate binding initiate the global conformational changes. Comparison of the binding pockets reveals that Y263 on TM7 and R305 on TM8 are associated with substrate binding and the conformational changes during the transport cycle (Fig.5B). In the IF and OF states, there is no interaction between Y263 and substrate in the binding pocket. In the OC states (state 4 and 8 in Fig.5B), however, the phenol hydroxyl group of Y263 hydrogen bonds with both R305 (TM8) guanidinium group and the substrate, thereby connecting the extensive hydrogen bond network among TM1, TM2, TM7, TM5, TM10, and TM11. The importance of Y263 and R305 is consistent with previous mutational studies.^22^ Y263F and R305K mutants in NarK completely abolished the transport activity even under high isopropyl-b-D-thiogalactoside (IPTG) concentration, which is used to induce protein expression. ^22^ Whereas R89K mutant rescued the 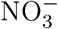-uptake activity under the highest IPTG concentration.^22^ These results support that the hydrogen bond between the side chains of Y263 and R305 is critical for the transport activity.

This hydrogen bond network rearrangement involving Y263 and R305 provides a plausible mechanism of the coupling between substrate binding and conformational changes (Fig.5B): In IF or OF states, the anionic substrate initially binds weakly to the binding site and neutralizes the basic residues: R89 and R305. At this stage, Y263 does not participate in binding. (2) The side chain of Y263 then moves closer toward and interacts with the substrate. This elicits the tighter substrate binding to the transporter. (3) The binding energy released by this stronger interaction is used to overcome the energy barrier from IF to OC or OF to OC transitions. The compact hydrogen bond network at the binding pocket turns on the helix bending motions. Since these helices constitute the N and C bundles, the bending motions directly result in the global conformational changes. Taken together, Y263 and R305 play a pivotal role in the coupling mechanism of the substrate binding and global conformational changes.

### A model for substrate recognition, binding, exchange, and translocation

The last question is which residues regulate substrate recognition, binding, and release? To identify key residues in substrate recognition, binding, and transport, we calculated interactions between NarK and substrates during the course of MD simulations. The contact frequency computation reveals that 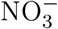 and 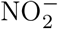 take the same transport pathway (central panel in Fig.6). The exchange between 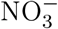 from the outside and 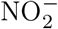 from the inside can be described by the following steps (Fig.6).

**Figure 6:**
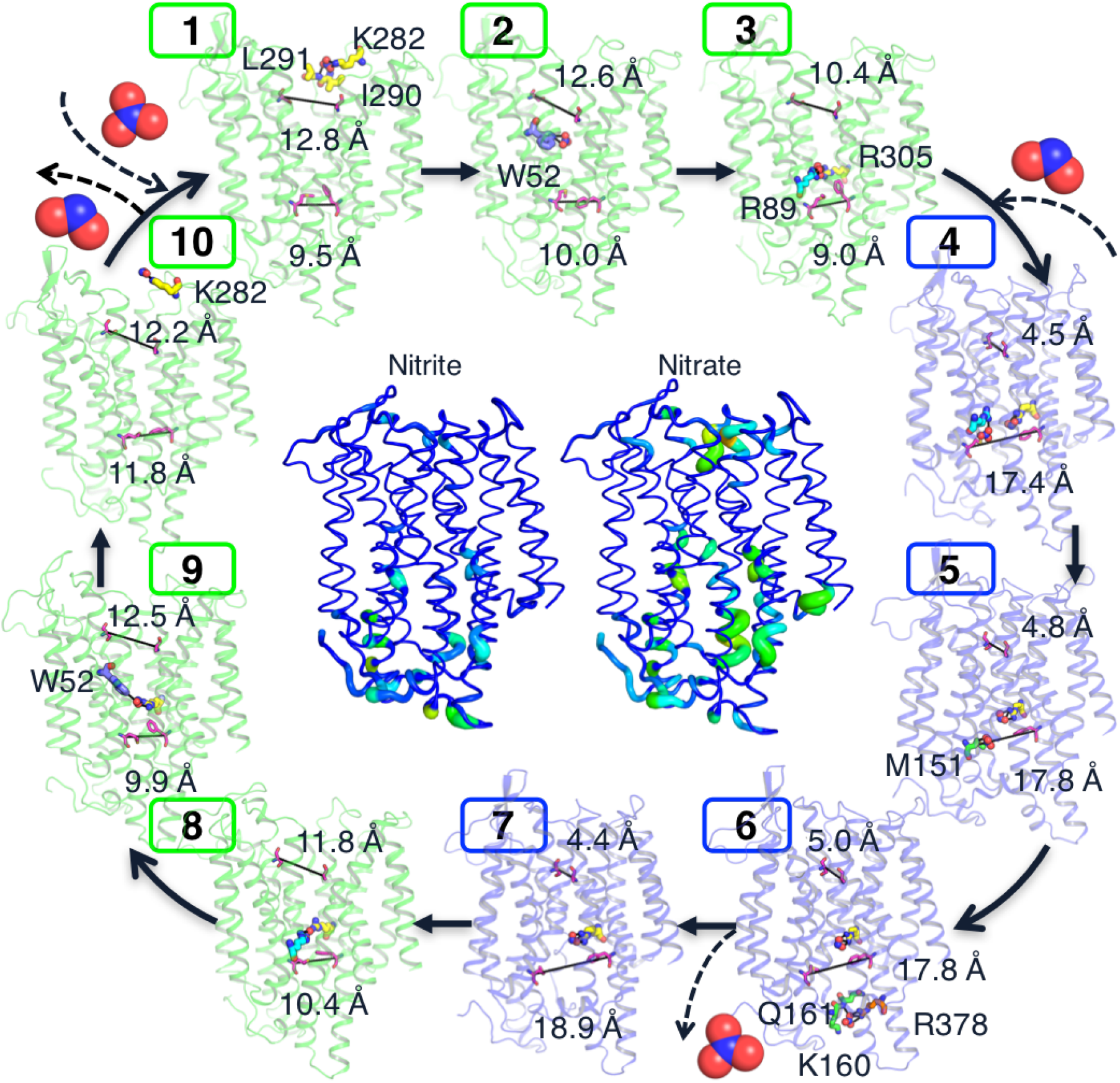
Substrate recognition, binding, exchange, and translocation. The central panel shows the MSM-weighed substrate-residue contact frequency for 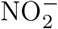 and 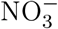. The tube thickness indicates the frequency of residue-substrate contact. (1-3) In the OF state, 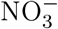 binds to NarK. (4) Structural transitions from OF to IF move R89 and R305 apart. 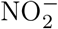 from inside solution binds the other unoccupied arginine. (5-7) 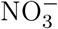 slides down to the cytosolic side. (8) The release of 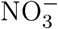 permits the closing of inside gates and triggers the conformational transitions from IF to OF. (9-10) 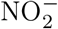 follows the same translocation pathway with 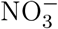 and gets released to the outside. For clarity, only R89 and R305 were shown in the binding site to illustrate the substrate binding. The dotted black lines represent the distances between gating residues (magenta sticks). Substates and relevant residues were represented as ball-and-sticks model.

(1) At first, the extracellular gating residues are ~12.8 Å apart and favor 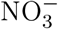 binding at the extracellular side. The positively charged residue K282 on the extracellular side of TM7 acts as a “hook", which appears to recruit the negatively charged 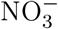 and escort the 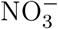 deep inwards to make contact with the backbones of L291 and I290 on TM8 (Fig.6(1)). (2) After transient coordination with K282 and nearby L291 and I290, 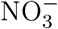 rapidly diffuses deep into the pore channel through W52 (Fig.6(2)). (3) 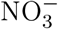 binds to the two positively charged and highly conserved residues (R89 and R305) in the binding site. 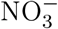 binding weakens the electrostatic repulsion between R89 and R305 and forms extensive hydrogen bond network in the binding pocket. The formation of hydrogen bonds between Y263, R305, and the substrate turns on the helix bending motions, which enables the outward halves of the core domains to approach each other and facilitates the conformational change from the OF to OC state. At this juncture, the extracellular gating distance decreases to ~10.4 Å (Fig.6(3)). (4) The formation of three layers of interactions finally closes the extracellular gate (~4.5 Å), which occurs with an opening of the intracellular vestibule (~17.4 Å). Due to the outward bending of TM5, TM10, and TM11, R89 and R305 move away from each other. 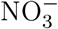 is now bound to only one of the arginine residues. This makes space for the binding of 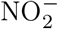 from the inside solution (Fig.6(4)). (5) The repulsion between 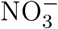 and 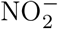 facilitates the dissociation of 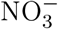 from the central binding site. 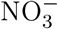 enters the intracellular part of NarK and forms a hydrogen bond with M151 (Fig.6(5)). (6) 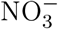 is released to the inside solution mediated by positively charged and polar residues (K160, R378, and Q161) located on the inward surface (Fig.6(6)). (7) Once 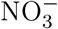 dissociates, only 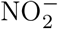 is bound in the binding site. This allows the closure of the inside gate and leads to conformational transition from IF to OC (Fig.6(7)). (8) As the closure of the inside gate (~10.4 Å) through the inward bending of TM5, TM10, and TM11, 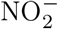 binds to both R89 and R305 which triggers the opening of the outside gate (~11.8 Å) (Fig.6(8)). (9) 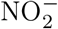 follows the same pathway as 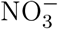 to get released to the outside solution. 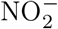 is dissociated from the binding site through W52 (Fig.6(9)). (10) 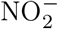 is released to the outside solution mediated by the positively charged K282 on the outward surface (Fig.6(10)).

## Discussion

In an attempt to decipher the molecular origin of MFS antiporters, we studied a bacterial 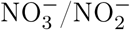 antiporter, NarK. Extensive all-atom MD simulations (~300 *μ*s in total) allow us to characterize the unbiased dynamics along the complete substrate exchange cycle of NarK. To establish NarK as a model for understanding MFS antiporters, we focus on addressing the following structural and mechanistic questions.

(1) Can antiporters be distinguished from uniporters and symporters according to protein architecture and conformational transition mechanism? Simulation results suggest that NarK adopts the canonical MFS fold and rocker-switch mechanism. Rather than the overall protein architecture, a few residues in the binding pocket and translocation pathway result in the different MFS functions.

(2) How is the empty transporter prevented from changing conformations in antiporters? In NarK, two absolutely conserved and positively charged arginine residues (R89 and R305) in the central binding site prevent the closure of the unbound protein. Substrate (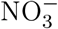 or 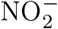) binding neutralizes the charge and weakens the electrostatic repulsion, thus permitting the conformational transitions. Paired basic residues were also found critical for substrate binding in several other MFS antiporters (*i.e.* GlpT, UhpT, and OxlT^53–57^). This suggests that paired basic residues in the binding pocket may be the common structural determinant for the exchange function in antiporters which transport anionic substrates.

(3) How does antiporter distinguish and switch two cargos? Breakage and formation of hydrogen bonds rearranges the binding pocket to differentiate and fit two different cargo molecules. The additional hydrogen bonds formed by the extra oxygen of 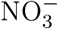 connect the two halves of NarK tighter than 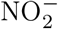 and result in higher binding affinity of 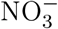. Regarding the switching mechanism, our results support a state-dependent substrate exchange mechanism. This is due to relative spacing between R89 and R305. Both substrates bind in the IF state and the electrostatic repulsion between 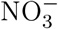 and 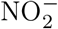 speeds up the release of one substrate. By contrast, R89 and R305 in OF state are closer to each other by ~2.0 Å and both make contacts with the substrate, leading to sequential binding.

(4) How is substrate binding coupled with global conformational changes? The coupling between substrate binding and global conformational changes is ensured through the movement of Y263 and R305 at the binding pocket. Upon substrate binding, the side chain of Y263 moves closer to interact with R305 and the substrate and forms an extensive hydrogen bond network. The shrinkage of the binding pocket triggers the helix bending motions and directly results in the global conformational changes.

(5) Which residues regulate substrate recognition, binding, and release? This work identifies all the relevant residues along the translocation pathway and explains the complete substrate recognition, binding, exchange, and release process.

Our results also identify structural features determining the high-affinity transport activity of all NNP proteins. The highly conserved glycine residues serve as the pivot points of helix bending, which is energetically efficient with a relatively low amount of energy (~2.5 ± 0.1 kcal/mol) required to complete the cycle from IF to OF states. Another structural feature is the gating residues lining up the translocation pathway. Previous studies of GlpT antiporter suggested that salt bridges are important for gating on both sides of the binding pocket.^59^ However, such salt bridges are lacking on both sides of the substrate-binding site of NarK. Two hydrophobic interaction layers (layer E3 and layer I1 in Fig.3) occlude the substrate and require only modest local structural changes to break or form.

Summarized, by performing the extensive all-atom MD simulations, the present study provides the complete picture of the NarK antiporter function (Fig.6). The detailed information provided in this study sheds light on the fundamental mechanism of all MFS antiporters. Furthermore, the relevant residues identified in this work can be used to engineer NNP proteins in crops to achieve higher crop productivity.

### Data and Code Availability

Source codes for adaptive MD simulations, MSMs hyper-parameters selection, and MSMs construction used in the paper are available at Github: https://github.com/ShuklaGroup/NarK_Structure_2021_Files. Detailed explanation of the procedures is reported in the Method Details section. All softwares and libraries used are reported in the Method Details section, together with the Key Resources Table.

## Method Details

### Molecular Dynamics (MD) Simulation

The 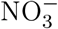-bound OC NarK crystal structure (PDB ID: 4U4W^22^) was downloaded from Protein Data Bank (PDB)^60^ and used as the starting coordinates for MD simulations. The chain termini were capped with neutral acetyl and methylamide groups. The membrane-protein MD system was built using the Membrane Builder plugin in CHARMM-GUI.^61^ The protein was embedded in a phosphatidylcholine (POPC) bilayer, solvated in a box with dimension of 76Å*76Å*102Å with TIP3P water molecules, and neutralized by adding 22 Na^+^ ions. 28 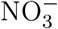 and 28 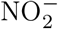 ions were randomly added to the MD system using Packmol.^62^ The final MD system contains 51,315 atoms. The MD system was first minimized using conjugate gradient method for 10,000 steps. Next, the system was slowly heated from 0 K to 10 K, and then 10 K to 300 K over a period of 1 ns each under canonical (NVT) condition. The system was further equilibrated in isothermal-isobaric (NPT) condition for 50 ns.^63^ All simulations were implemented using Amber14 package^64^ employing Amber ff14SB^65^ and GAFF^66^ force field, and carried out in the NPT conditions (300K and 1 atm) maintained using a Berendsen thermostat and a Berendsen barostat.^67^ Periodic boundary conditions (PBC) were applied to all simulations, Particle Mesh Ewald was used to treat long-range non-bonded interactions, ^68^ and the SHAKE algorithm was used to constrain hydrogen-containing bonds. ^69^ A 2 fs time step was used throughout all simulations.

### Adaptive Sampling

To overcome the timescale gap between the large timescale for the 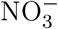 and 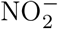 exchange processes in NarK and the short range timescale that can be explored in MD simulations, we employed a Markov state model (MSM) based adaptive sampling protocol. Adaptive sampling is a widely used sampling method and has demonstrated the efficiency of capturing various biologically relevant processes and tremendous value in time and resource savings. ^70–72^ The adaptive sampling scheme used in this work is based on the counts of different states in order to discover new states quickly. The procedure of adaptive sampling is:

1. Run a series of short MD simulations from initial seeds in parallel. For the first round, 250 parallel simulations were launched from the crystal structure (PDB ID: 4U4W^22^). After the first round, around 200 parallel simulations were launched per round. The simulation length is set as 30 ns to both satisfy the Markovian assumption and maximize the sampling efficiency.
2. Cluster all the simulation data collected so far using the K-means algorithm with number of clusters specified as 5000. Two biologically relevant metrics are selected as clustering metrics: extracellular gating distance (Ser56C*α* (TM1) - Ala275C*α* (TM7)) and intracellular gating distance (Met151C*α* (TM4) - Phe370C*α* (TM10)).
3. Select around 200 structures from the least populated clusters as seeds for the next run of simulation.
4. Repeat steps 1-3 until the sampling reaches the convergence criterion: the free energy landscapes stop varying as rounds continue.
5. Build a Markov state model (see the Markov state model (MSM) Construction section in STAR Methods for details) from the final data set to capture the proper thermodynamics and kinetics. MSMs will help correct any sampling bias introduced by selecting the starting conformations from each round of simulations according to the least populated clusters instead of a Boltzmann distribution.

A total of ~300 *μ*s simulation data were obtained and used for further data analysis. The in-house python codes used to perform adaptive MD sampling are available at https://github.com/ShuklaGroup/NarK_Structure_2021_Files.

### Markov state model (MSM) Construction

An issue with adaptive sampling is the introduced sampling bias as each new round of simulations starts from the least populated states, which may alter the real equilibrium population of the states. To eliminate the sampling bias, we constructed Markov state model (MSM) to statistically stitch all the short simulation data and estimate the transition probability matrix between all of the conformational states.^32,73^ The procedure of constructing a MSM is: (1) featurizing the trajectory data using a set of C*α* distances between residues; (2) decomposing the featurized data using the time-lagged independent component analysis (tICA) technique, which finds the slowest collective motions in the system through linear combinations of the input features (C*α* distances between residues in our case); and (3) clustering the decomposed data into conformational microstates using Mini-batch K-means clustering algorithm.

To select the optimal hyper-parameters (C*α* contacts, number of tICA components, and number of clusters) systematically and automatically, we employed a genetic algorithm based technique developed from our lab^38^ (Supplementary Table 1 and 2). The source codes and the resulting data associated with this algorithm are available at https://github.com/ShuklaGroup/NarK_Structure_2021_Files. The whole idea is mimicking the natural selection and evolving the best combinations of these hyper-parameters based on the fitness score. The generalized matrix Raleigh quotient (GMRQ) is used as the fitness score to quantify the quality of MSM models, as GMRQ is the sum of the eigenvalues of the transition matrix estimated from MSM and the higher the GRMQ, the better the MSM is at capturing the slowest motions in the system.^74,75^ The workflow for the genetic algorithm based search of optimal hyper-parameters consist of:

1. Prepare the input file which specifies four genetic algorithm parameters: *N_ITERATIONS*, *populationSize*, *percentMutation*, and *percentCrossover*. The input file used in this work *nark.inp* is available at https://github.com/ShuklaGroup/NarK_Structure_2021_Files.
2. Generate a pool of all possible residue pairs to explore. A total of L(L-1)/2 residue pairs were obtained for NarK where L = 447 is the protein length. The generated file *compatiblePairs.txt* is available at https://github.com/ShuklaGroup/NarK_Structure_2021_Files.
3. Run the genetic algorithm which automatically constructs MSMs for different combinations of hyper-parameters, evaluates the quality of MSMs with GMRQ, and generates the new generation according to the GMRQ. The algorithm stops when it reaches maximum number of iterations (*N_ITERATIONS* = 40) specified in the input file. The output files for all the 40 iterations (*iter_*_output_sets.txt*) are available at https://github.com/ShuklaGroup/NarK_Structure_2021_Files.

The convergence of GMRQ scores is shown in Supplementary Fig.6A. Eventually, the highest scored combination of hyper-parameters, that is 61 C*α* contact distances, 10 tICA components, and 400 clusters, was chosen for the final MSM construction (details are shown in Supplementary Table 3). A Markovian lag time 15 ns was chosen from the implied timescale plot to construct the MSM (Supplementary Fig.6B). In-house python codes were used to construct the final MSM model (available at https://github.com/ShuklaGroup/NarK_Structure_2021_Files).

The Osprey variational cross-validation package was used in the genetic algorithm workflow to cross validate the MSMs by varying the hyper-parameters.^76^ MSMBuilder 3.6^77^ was used to construct MSMs, CPPTRAJ^78^ module in AMBER14^64^ and MDTraj1.7^74^ were used to analyze simulation trajectories, and VMD1.9.2^79^ and PyMol 3^49^ were used to visualize MD snapshots.

## Quantification and Statistical Analysis

Data analysis were performed using MDTraj1.7,^74^ MSMBuilder 3.6,^77^ PyEMMA2.5.7,^41^ along with in-house python codes (available at https://github.com/ShuklaGroup/NarK_Structure_2021_Files). Plot graphics were generated with matplotlib.^80^

To quantify the uncertainties of the MSM model, we estimated a Bayesian MSM using PyEMMA2.5.7.^41^ Bayesian MSMs construct a sample of reversible transition matrices using a Gibbs sampling scheme and are therefore commonly used to quantify the statistical uncertainties for all observables derived from MSM.^39,40^ In this work, we estimated the Bayesian MSM with 100 samples and 95% confidence interval. The sample mean of the stationary distribution was then used to reweigh the free energy landscapes (Supplementary Fig.4). In order to estimate the uncertainties of free energies, we computed the free energy differences between the Bayesian MSM reweighted landscapes and the maximum likelihood MSM reweighted landscapes (Supplementary Fig.6). The maximum free energy deviation is less than 0.1 kcal/mol. We therefore use 0.1 kcal/mol with 95% confidence as the upper bound for the approximation errors made by modeling protein dynamics with a MSM.

A caveat for the free energy analysis is that Berendsen thermostat was used to control temperature, which may affect the distributions of the desired ensemble. ^81,82^ To quantify the errors introduced by the thermostat, we re-ran all the simulations using Langevin thermostat with collision frequency of 2 ps^*−*1^.^83,84^ The protocol of the new adaptive MD sampling is: (1) initialize ~ 10, 000 trajectories from 400 MSM states (~ 25 trajectories per MSM state), (2) cluster all the simulation data using the K-means algorithm with number of clusters specified as 5000. Two biologically relevant metrics are selected as clustering metrics: extracellular gating distance (Ser56C*α* (TM1) - Ala275C*α* (TM7)) and intracellular gating distance (Met151C*α* (TM4) - Phe370C*α* (TM10)), (3) select around 500 structures from the least populated clusters as seeds for the second run of simulation. An aggregate simulation time of ~230 *μs* was obtained. We reconstructed a MSM for the new simulation data and generated new free energy landscapes for comparison (Supplementary Fig.4). The comparison of all four free energy landscapes shows that (1) the overall shapes of free energy landscapes do not vary, (2) the free energy differences between different states remain the same, (3) although there are minor free energy differences for some regions, the differences are much less than 0.1 kcal/mol which is the error bar of free energies determined through uncertainty test. Therefore, we suggest that the free energy analysis is still valid, because the errors caused by the thermostat choice are much less than the error bars.

## Acknowledgement

Authors thank the Blue Waters sustained-petascale computing project, which is supported by the National Science Foundation (Awards OCI-0725070 and ACI-1238993) and the state of Illinois. D.S. acknowledges support from the New Innovator Award from the Foundation for Food and Agriculture Research and NSF Early Career Award (MCB 1845606). J.F. was supported by Chia-chen Chu Fellowship and Harry G. Drickamer Graduate Research Fellowship from University of Illinois, Urbana-Champaign, IL.

## Author Contributions

D.S. conceived and supervised this study. J.F. and B.S. performed simulations. J.F. analyzed the simulation data, made figures, and wrote the manuscript with inputs from D.S.

