## Supplementary_Information for "How antiporters exchange substrates across the cell membrane? An atomic-level description of the complete exchange cycle in NarK"

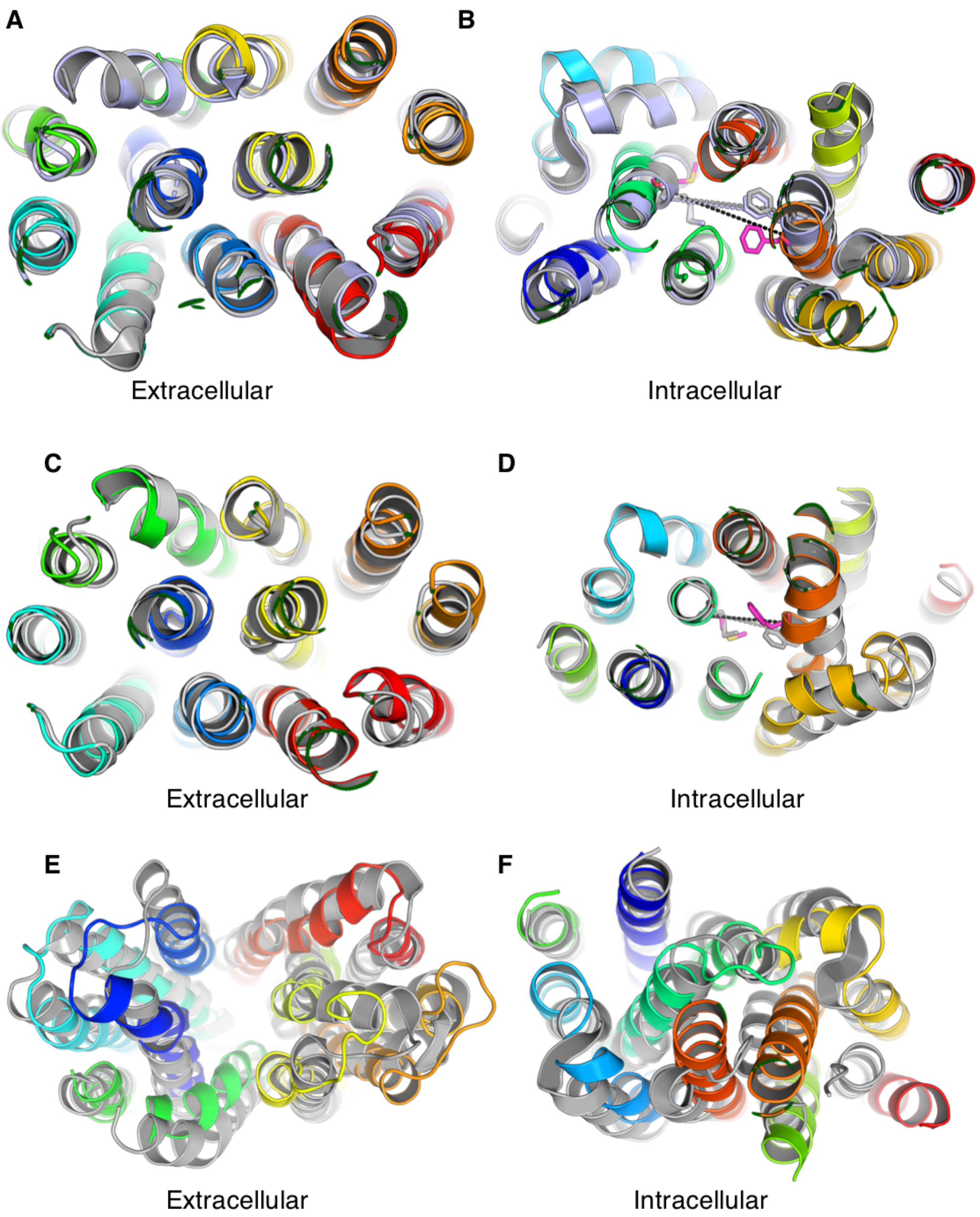


Figure S1: **Related to Figure 1; Overlay of the predicted structures with crystal structures.** (A-B) Overlay of the predicted IF structure with *apo* IF (PDB ID: 4U4V; light blue) and $NO_3^-$-bound IF (PDB ID: 4U4T; gray) crystal structures. (A) Extracellular view. (B) Intracellular view highlighting the difference in the intracellular gating distance (Met151C$\alpha$ - Phe370C$\alpha$). The intracellular distance is 15.2 Å, 12.9 Å, 13.4 Å for the predicted IF, 4U4V, and 4U4T. (C-D) Overlay of the predicted OC structure with $NO_3^-$-bound OC crystal structure (PDB ID: 4U4W; gray). (C) Extracellular view. (D) Intracellular view highlighting the difference in the intracellular gating distance (Met151C$\alpha$ - Phe370C$\alpha$). The intracellular distance is 8.6 Å and 11.5 Å for the predicted OC and 4U4W. (E-F) Overlay of the predicted OF structure with the OF structure of a fucose transporter (FucP, PDB ID: 3O7Q; gray). (E) Extracellular view. (F) Intracellular view.

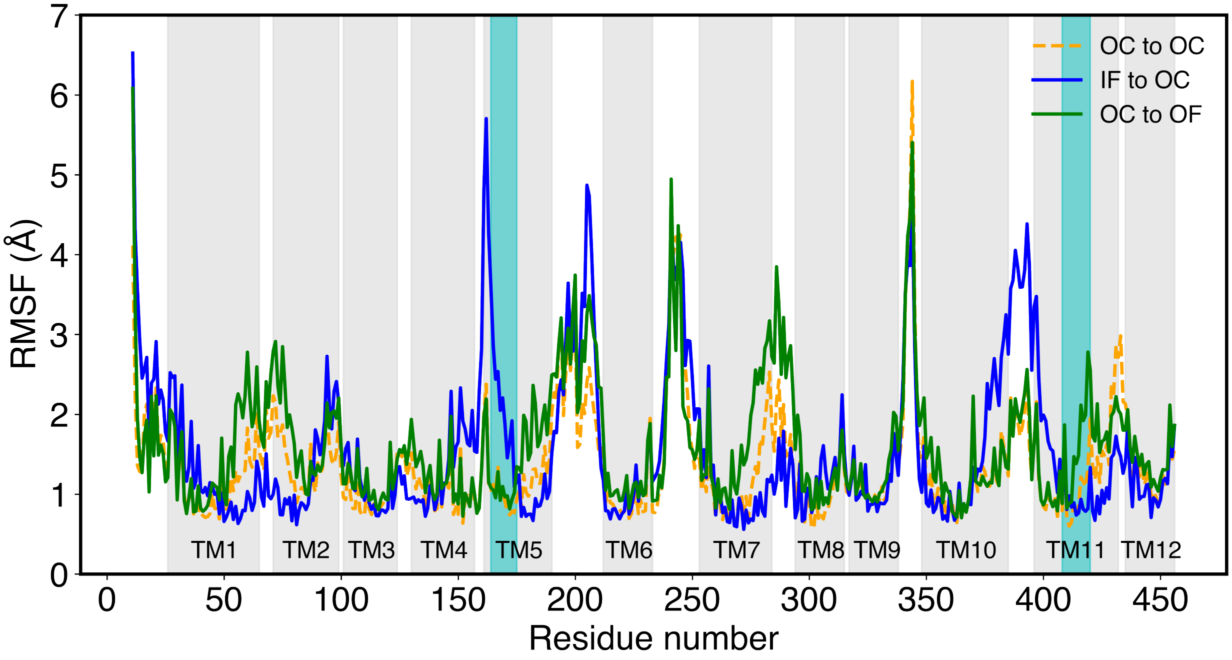


Figure S2: **Related to Figure 2; Root-mean-square fluctuation (RMSF) of each C$\alpha$ atom in NarK during the IF to OC and the OC to OF transition.** The cyan regions represent two $NO_3^-$ signature motifs in TM5 and TM11.

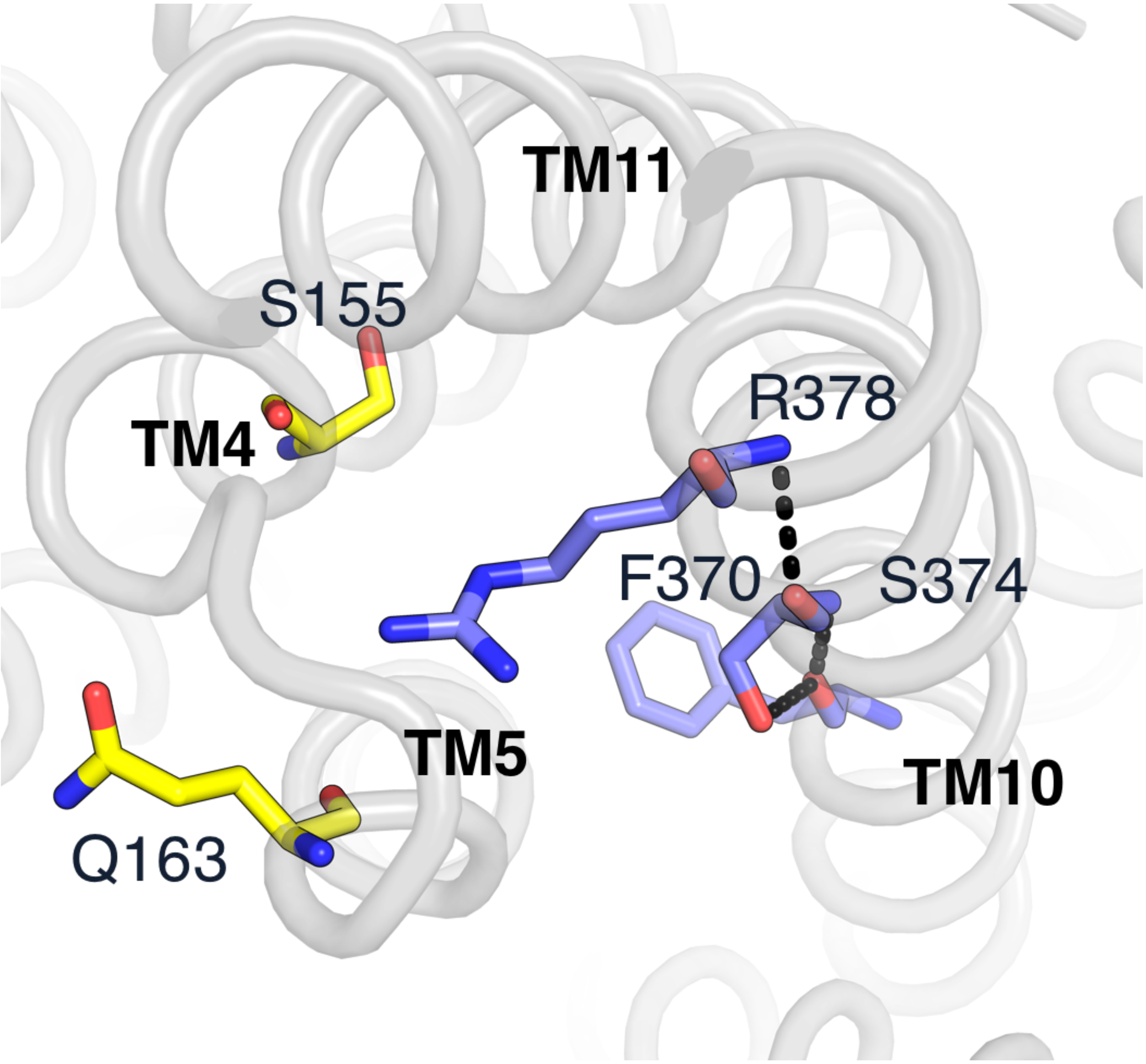


Figure S3: **Related to Figure 3; The missing intracellular interaction layer 2 in the $NO_3^-$-bound OC crystal structure (PDB ID: 4U4W; gray).** The hydrogen bond network is missing due to the unfavorable position of Q163 side chain.

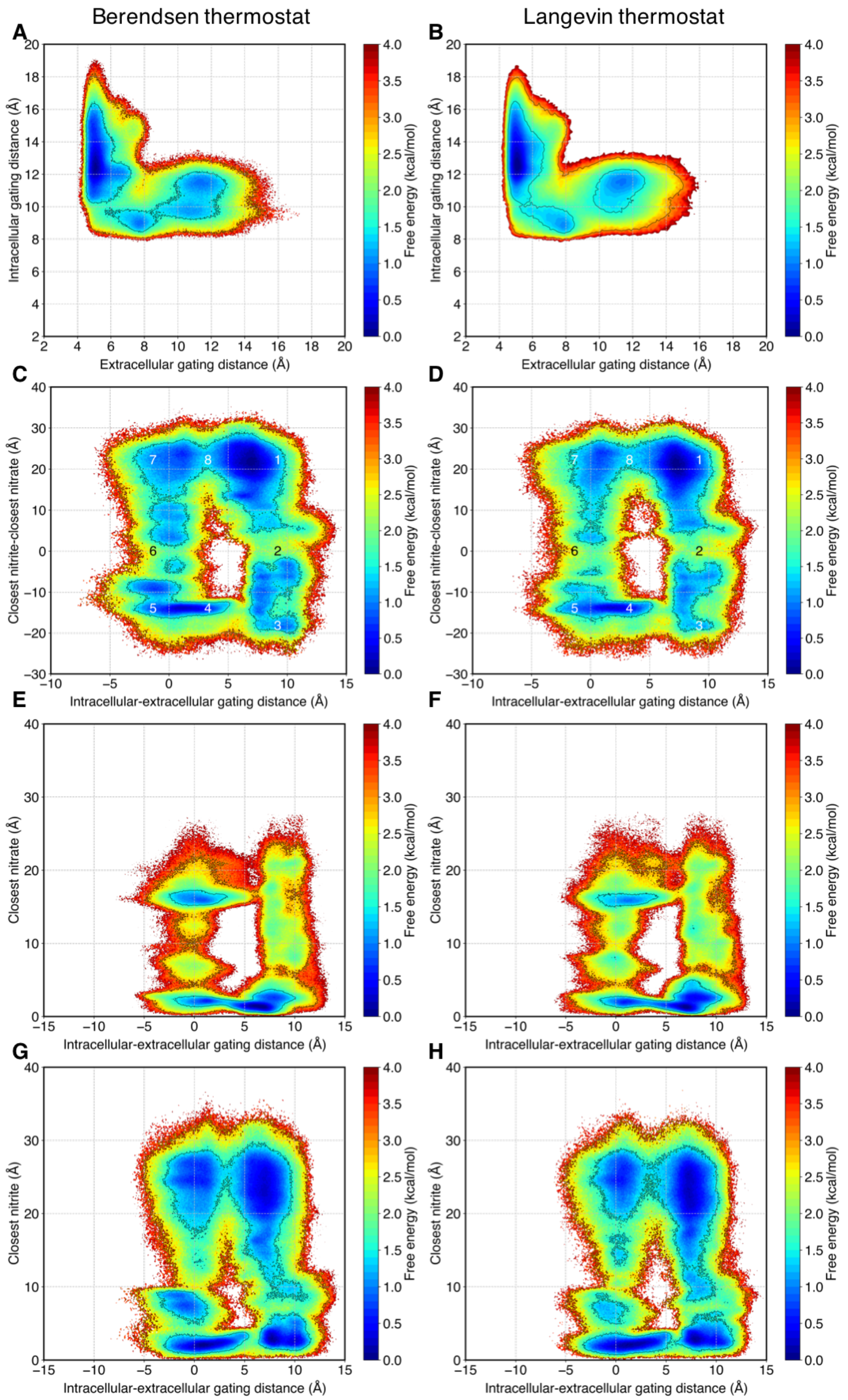


Figure S4: **Related to STAR Methods, Figure 1, and Figure 4; Comparison of MSM-weighted free energy landscapes between Berendsen (left column: A, C, E, G) and Langevin thermostat (right column: B, D, F, H).** MSM-weighted free energy landscapes of (A-B) the alternating access cycle, (C-D) the entire $NO_3^-/NO_2^-$ exchange cycle, (E-F) $NO_3^-$ translocation, (G-H) $NO_2^-$ translocation. The conformational states were depicted in C, D as (1) $NO_3^-$-bound IF, (2) *"apo"* IF, (3) $NO_2^-$-bound IF, (4) $NO_2^-$-bound OC, (5) $NO_2^-$-bound OF, (6) *"apo"* OF, (7) $NO_3^-$-bound OF, and (8) $NO_3^-$-bound OC.

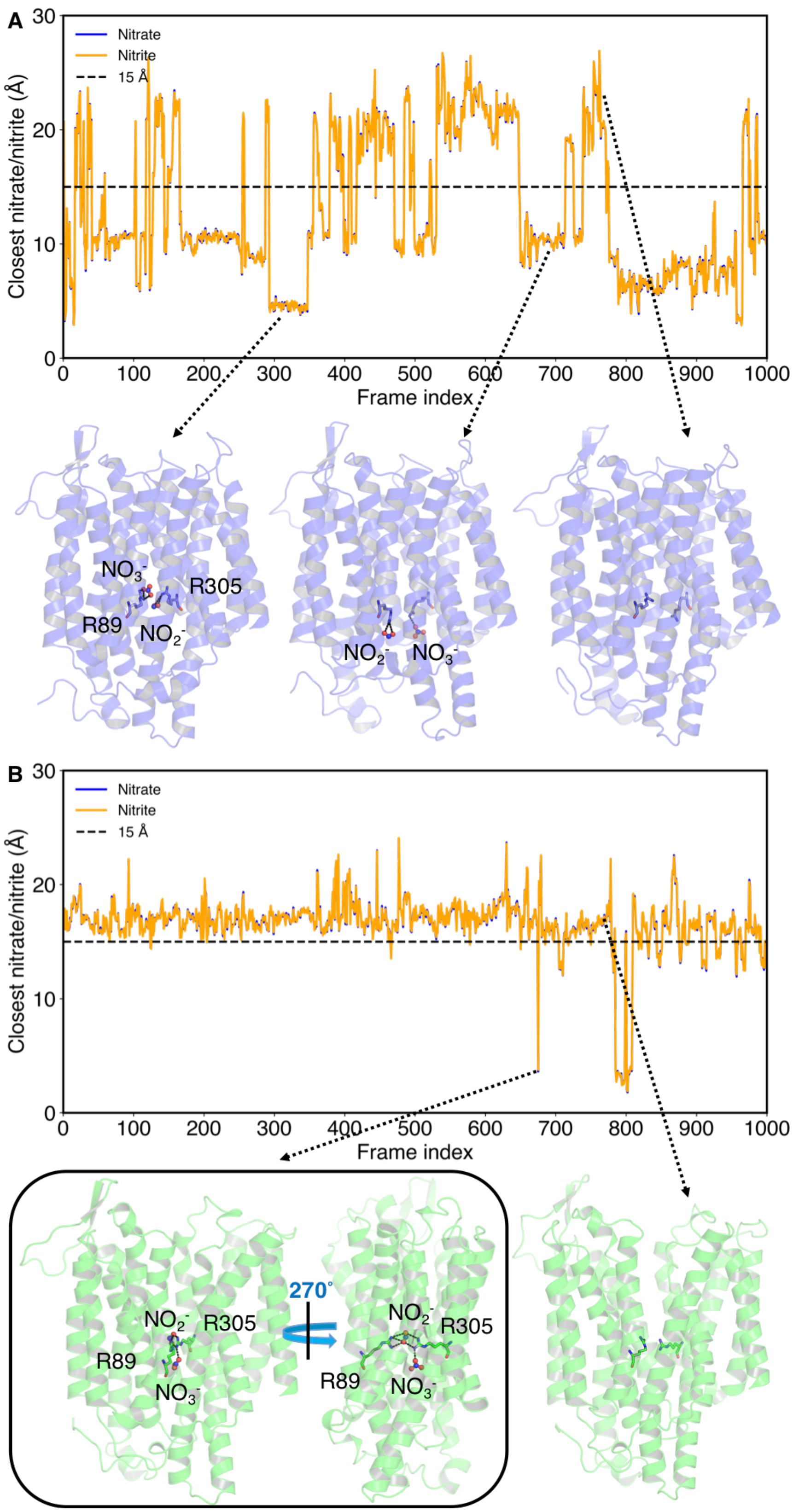


Figure S5: **Related to Figure 5; *Apo* versus both substrate binding in (A) IF and (B) OF states.** 1,000 MD snapshots were randomly chosen for each state. Representative snapshots for different distance conditions were visualized.

**Table S1: Related to STAR Methods; MSM lagtime estimation. All dihedral angles were used for featurization to gain initial estimates of MSM lagtime. 20 MSM models with varying number of tICA components and number of clusters were constructed. 15 ns was selected as an initial estimate of MSM lagtime based on the 20 implied timescale plots generated from 20 different MSM models.**

| **Featurization** | | |
|---|---|---|
| Dihedral | types=[‘phi’, ‘psi’], sincos=True | |
| **Decomposition** | **Components** | **lagtime (ns)** |
| tICA | 5, 10, 15, 20 | 0.1 |
| **Clustering** | **Clusters** | |
| Mini-batch k-means | 200, 400, 600, 800, 1000 | |
| **Model fitting** | **lagtime (ns)** | **Timescales** |
| MSM | 0.5 to 22 | 10 |

**Table S2: Related to STAR Methods; Osprey search range. The search range of MSM parameters for the genetic algorithm technique.**

| **Featurization** | | |
|---|---|---|
| C$\alpha$ contacts | | |
| **Decomposition** | **Components** | **lagtime (ns)** |
| tICA | 2, 5, 10, 15 | 0.1 |
| **Clustering** | **Clusters** | |
| Mini-batch k-means | 200, 400, 500, 600, 800, 1000 | |
| **Model fitting** | **lagtime (ns)** | **Reversiblity method** |
| MSM | 15 | MLE |
| **Scoring** | **Timescales** | |
| GMRQ | 5 | |
| **Cross-validation** | **Iterations** | **Test set size** |
| Shuffle & split | 5 | 0.5 |

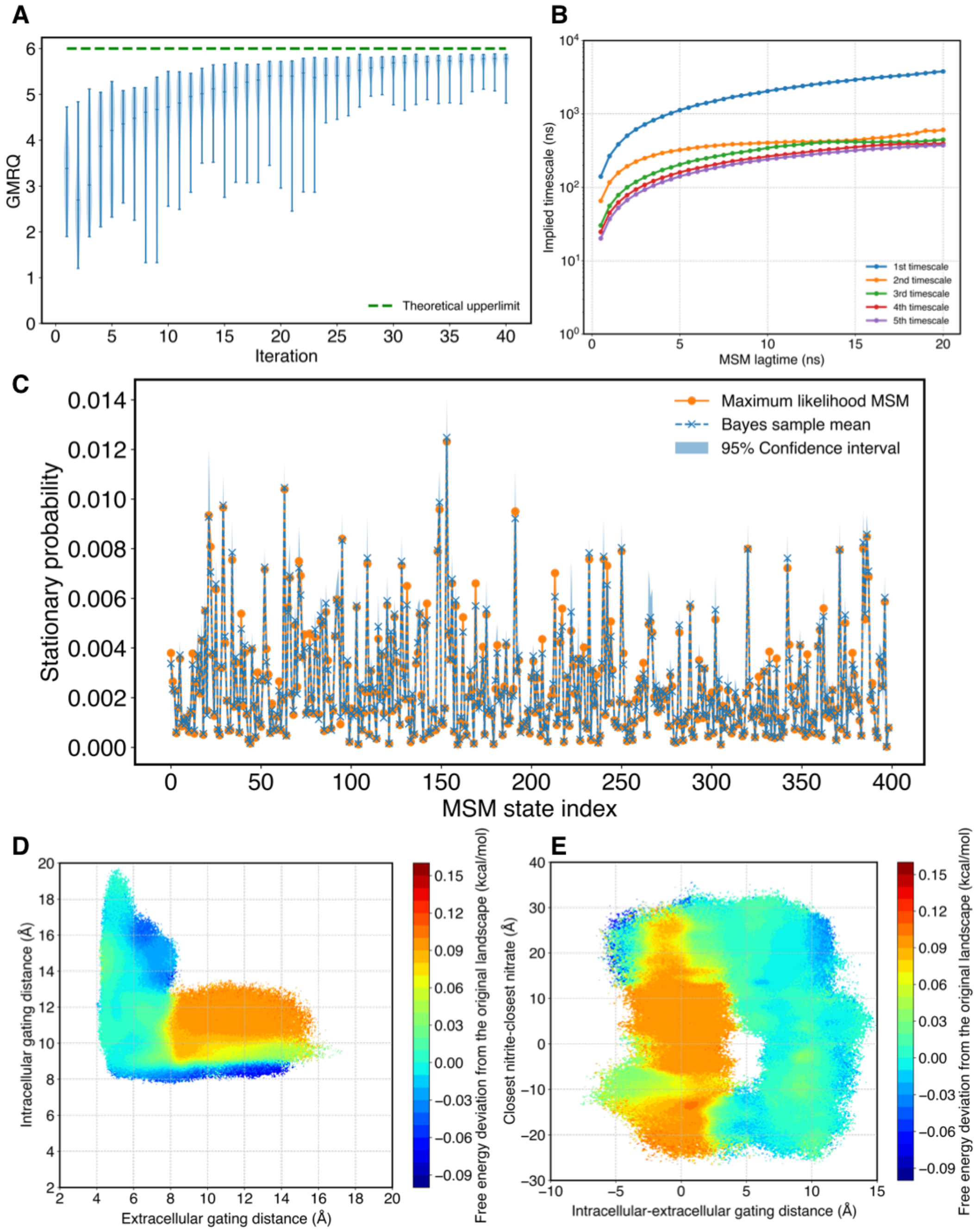


Figure S6: **Related to STAR Methods, Figure 1, and Figure 4; MSM construction and uncertainty quantifications.** (A) The convergence of generalized matrix Raleigh quotient (GRMQ) score. The green dashed line represents the theoretical upperlimit of GMRQ based on five timescales (GRMQ = 6). (B) Implied timescale plot for five slowest eigenvectors. The lagtime of 15 ns was chosen to build MSM to ensure the Markovian property. (C) Stationary probability of MSM states. The solid orange line refers to the maximum likelihood result; the dashed blue line and the shaded areas indicate sample mean and 95% confidence intervals computed using a Bayesian MSM with 100 samples. (D, E) The free energy error quantifications using the Bayesian MSM corresponding to Figure 1 (D) and Figure 4 (E).

**Table S3: Related to STAR Methods; Optimized MSM construction. The final set of parameters were chosen by the genetic algorithm technique.**

| **Featurization** | | | |
|---|---|---|---|
| C$\alpha$ contacts | Number of contacts = 61 | | |
| **Decomposition** | **Components** | **lagtime (ns)** | |
| tICA | 10 | 0.1 | |
| **Clustering** | **Clusters** | | |
| Mini-batch k-means | 400 | | |
| **Model fitting** | **lagtime (ns)** | **Reversiblity method** | **Population** |
| MSM | 15 | MLE | 100% |
| **Scoring** | **Timescales** | **Score** | |
| GMRQ | 5 | 5.88 | |
